# Herbicidal activity of fluoroquinolone derivatives

**DOI:** 10.1101/2021.06.26.450056

**Authors:** Michael D. Wallace, Aleksandra W. Debowski, Kirill V. Sukhoverkov, Joshua S. Mylne, Keith A. Stubbs

## Abstract

Development of herbicides with novel modes of action are crucial for weed control and to hinder herbicide resistance. An attractive novel herbicidal target is plant DNA gyrase, which has been demonstrated to be effectively inhibited by the known antimicrobial ciprofloxacin. Despite this good herbicidal activity ciprofloxacin is not suitable as a herbicide due to its antimicrobial activity, therefore, a diverse library of analogues was analysed to gain insight into the aspects required for herbicidal activity. This analysis revealed significant structural modifications were tolerated and that the fluoride at C-6 and a cyclic amino group at C-7 were not crucial for herbicidal activity. The analysis also revealed that these modifications also affected the antibacterial activity with one compound demonstrating good herbicidal activity and weak antibacterial activity, against both Gram-positive and Gram-negative bacteria.

## Introduction

In the global production of food and plant-based products, herbicides are vital for effective management of weeds.^1^ However, since their introduction, resistance to herbicides is becoming more prevalent. To date, 263 weed species have populations resistant to one or more herbicide chemistries.^2^ To combat this, an important strategy is to introduce new herbicides, specifically with new modes of action.^3^ This will ultimately help to diversify herbicide based weed control strategies, hindering the emergence of resistance. Of significant concern is that until the very recent release of tetflupyrolimet and cyclopyrimorate, developed by FMC Agricultural Solutions and Mitsui Chemicals Agro, respectively,^4^ it was over 30 years since the last novel mode of action herbicide. Thus, introducing compounds like these must continue regularly for effective control of herbicide resistance.

Recently the compound ciprofloxacin **1** (Figure 1) was found to have good herbicidal activity against the model plant *Arabidopsis thaliana* via its target DNA gyrase.^5^ This enzyme was originally characterised in *Escherichia coli*^*6*^ and is an essential type II topoisomerase that relieves DNA supercoiling.^7^ As DNA gyrase is essential to bacteria and not present in mammals, it is an ideal antibiotic target, with fluoroquinolones a successfully developed class of antibiotics. Plant DNA gyrase was characterised in the 2000s and is similarly essential for plant growth, making it an attractive target for herbicide research.^8-10^

**Figure 1.**
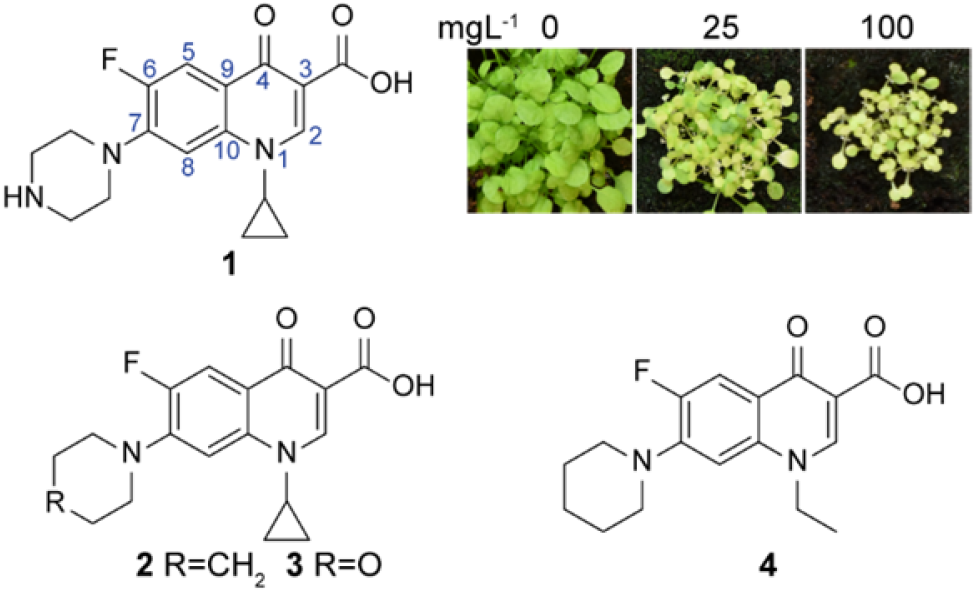
Previous studies into the structure and herbicidal activity of ciprofloxacin **1**, and the structures of previously prepared analogues **2**-**4**.

As DNA gyrase is essential for both plants and bacteria, any herbicide to have this mode of action needs to have negligible antibacterial activity. A previous study used **1** as a lead compound, making minor structural changes at the N-1 and C-7 positions in attempts to maintain herbicidal activity but lose antibacterial activity.^11^ Some success was had with **2** and **3** (Figure 1) that had a substitution of the secondary nitrogen of the piperazine ring with a methylene group or an oxygen, respectively. Analogues **2** and **3** had a 1.7- and 2.5-fold increase in herbicidal activity compared to **1**, respectively but a 128- and 32-fold reduction in activity against *E. coli*. The greatest selectivity shift was seen with **4**, which had a 1.7-fold reduction in herbicidal activity compared to **1**, but a 1000-fold decrease in activity against Gram-negative bacteria. Using *in vitro* supercoiling assays with recombinant plant and bacterial DNA gyrase, it was demonstrated that the results were not due to different affinity between DNA gyrase and DNA but rather the availability of the compound via plant or bacterial uptake. To expand upon these findings, we report here the herbicidal activity of a larger and more diverse library of **1** analogues, including modification of the core structure (Figure 2), to gain insight into the parts required for herbicidal activity.

**Figure 2.**
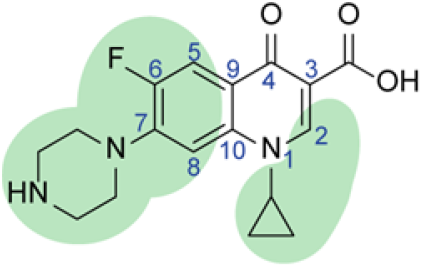
Ciprofloxacin **1** with areas of modification highlighted.

## Materials and Methods

### Ciprofloxacin analogues

Compounds **1, 5, 16** and **33** were purchased from Sigma Aldrich. Compounds **6**-**15, 17**-**32** and **34**-**64** were provided by Prof. Ulrich Jordis, Vienna University of Technology. The following analogues are DNA gyrase-inhibiting clinical antibiotics: ciprofloxacin **1**, danofloxacin **5**, levofloxacin **16**, oxolinic acid **19**, nalidixic acid **62** and trovafloxacin **64**.

### Physiochemical data

The calculation of physiochemical data and cluster analysis for the analogues were done as previously described.^12, 13^

### Screening for herbicidal activity

Approximately 30 seeds of *Arabidopsis thaliana* (wild type Col-0) were sown in a group on a pot (6 × 6 cm) of wet soil (Seedling Substrate Plus+, Bord Na Móna), and held for four days at 4 °C in a dark room to synchronise germination. Pots were then placed in growing conditions at 22 °C, 60% relative humidity in 16 h light/8 h dark conditions, for 18 days. All compounds were applied post-emergence, three- and six-days post germination. Compounds were dissolved in DMSO, at 10 g L^-1^ and diluted to 50 and 200 mg L^-1^ with a 0.02% surfactant solution, and total final DMSO concentration of 2%. The surfactant used was Brushwet (SST Australia, active constituent: 1020 g/L polyether modified polysiloxane). A 2% DMSO solution with surfactant was used as the negative control. Seedlings were treated with 500 μL of the solutions on each treatment day. Photos were taken on the final growth day and the images analysed for activity.

### Measuring herbicidal activity of active compounds

Seedlings were prepared and grown as described above and treated with 500 μL of active compounds at concentrations of 0, 3.125, 6.25, 12.5, 25, 50, 100 or 200 mg L^-1^ for **1, 5, 16, 19** and **39**, and of 0, 12.5, 25, 50, 100, 200, 400, 800 mg L^-1^ for **29**. All solutions contained 0.02% Brushwet and 2% DMSO. Photos were taken and images were analysed for growth and health of the *A. thaliana* plants, using ImageJ according to the methods of Corral *et al*.,^14^ with each compound tested in triplicate. Using this data IC_50_ values were calculated using GraphPad Prism 9.

### Antibacterial activity of herbicidal compounds

The Minimum Inhibitory Concentrations (MIC) against *Escherichia coli* (K12), *Pseudomonas aeruginosa* (ATCC 19429) and *Staphylococcus aureus* (ATCC 25923) were determined as previously described.^11^

## Results and Discussion

### Physiochemical analysis of analogues

The physiochemical data of the compounds were compared to commercial herbicides using the database available from Gandy *et al*.^12^ with an updated version from Sukhoverkov *et al*.^13^ From the cluster analysis performed (Figure 3) it was seen that for most properties (molar mass, partition coeffecient, distribution coefficient and polar surface area) the analogues generally fit within the bounds of common properties of herbicides. However, for the proportion of aromatic atoms, the majority of the analogues, including **1**, had percentages higher than what is typical for herbicides (Figure 3F). In addition, a large proportion of the analogues had reletively higher aqueous solubilities (Figure 3B and 3E).

**Figure 3.**
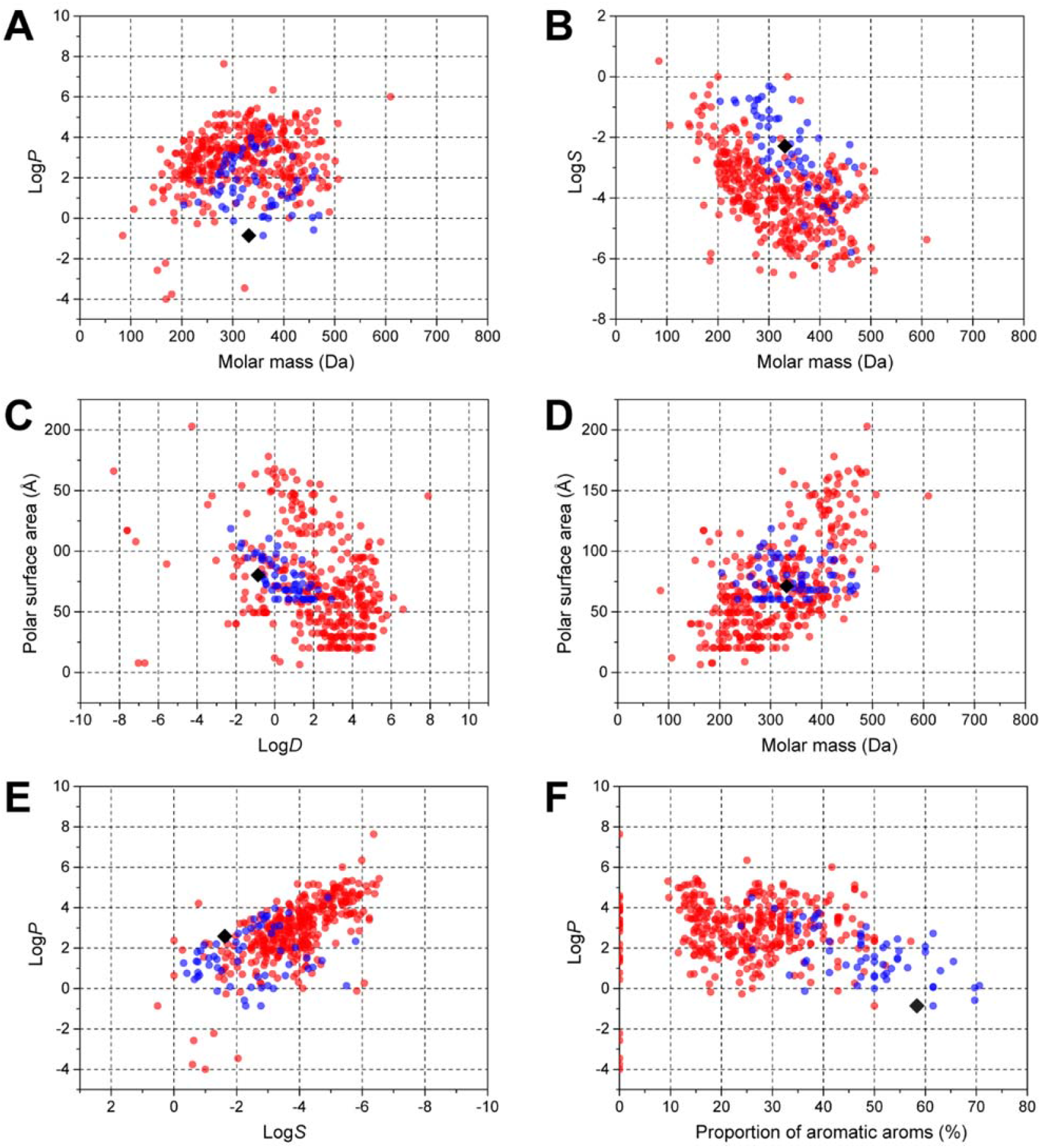
Cluster analysis of physiochemical properties of herbicides and ciprofloxacin analogues. (**A**) Molar mass and partition coefficient (Log*P*); (**B**) molar mass and aqueous solubility (Log*S*); (**C**) distribution coefficient (Log*D*) and polar surface area; (**D**) molar mass and polar surface area; (**E**) aqueous solubility (Log*S*) and partition coefficient (Log*P*); (**F**) proportion of aromatic atoms and partition coefficient (Log*P*). The black diamond represents **1**, the blue circles represent the 59 analogues of **1**, and red circles represent 360 commercial herbicides.^13^

### Herbicidal screening of ciprofloxacin analogues

To gain initial insight into the herbicidal potency of the library of compounds, they were initially screened for activity against *A. thaliana* at concentrations of 50 and 200 mg L^-1^ (Figure 4). Slight modification of the piperazinyl ring at C-7 of **1** with a bicyclic motif as in **5** resulted in the retention of activity. Fluoroquinolone derivatives **6**-**15** with various C-7 amino groups lacked activity. Levofloxacin **16**, which has a fused tricyclic core, had herbicidal activity at 50 mg L^-1^. However, analogues of ofloxacin (racemate of **16** and its enantiomer) **17** and **18** had no activity. The linear tricyclic oxolinic acid **19** was also active at 50 mg L^-1^, despite lacking the C-6 fluoride or a C-7 cyclic amine seemingly crucial for activity of many analogues tested here and previously.^11^ The compounds **20**-**28**, which are analogues of **19** with varied heterocyclic 5- and 6-member rings replacing the methylenedioxy group, all lacked herbicidal activity. The derivative **29** interestingly has a 4-fluorophenyl substituent at N-1 and a 7-ethoxy group, but modest activity only at 200 mg L^-1^. The 7-alkyl **30** and 7-thiol **31** and **32** linked derivatives had no activity. Structures with the quinolone-carboxylic acid core structure, and either H-, F- or Cl-substitution of the 6, 7 and 8 positions, showed no herbicidal activity for the *N*-cyclopropyl **33**, -ethyl **34** and **35**, -4-fluorophenyl **36**, -2,4-difluorophenyl **37** and -aminomethyl **38**. However, the derivative **39** with a thiazolidinyl group across the N-1 and C-2 positions, had herbicidal activity at 50 mg L^-1^. Like **19, 39** lacks a cyclic amine at C-7, further suggesting this group might not be as crucial as previous results suggested.^11^ The fused tetracyclic derivatives **40**-**42** lacked activity, as well did the tricyclic benzoquinolone derivatives **43**-**50**. The benzothieno derivatives **51**-**60** also showed no herbicidal activity. Nalidixic acid **61**, its precursor **62** and trovafloxacin **63** had no observed herbicidal activity. This could be due to these derivatives being based off a 1,8-naphthyridine core, rather than a quinoline core. Notably, **61** and **63** are the first DNA gyrase inhibiting clinical antibiotics that lack herbicidal activity against *A. thaliana*.

**Figure 4.**
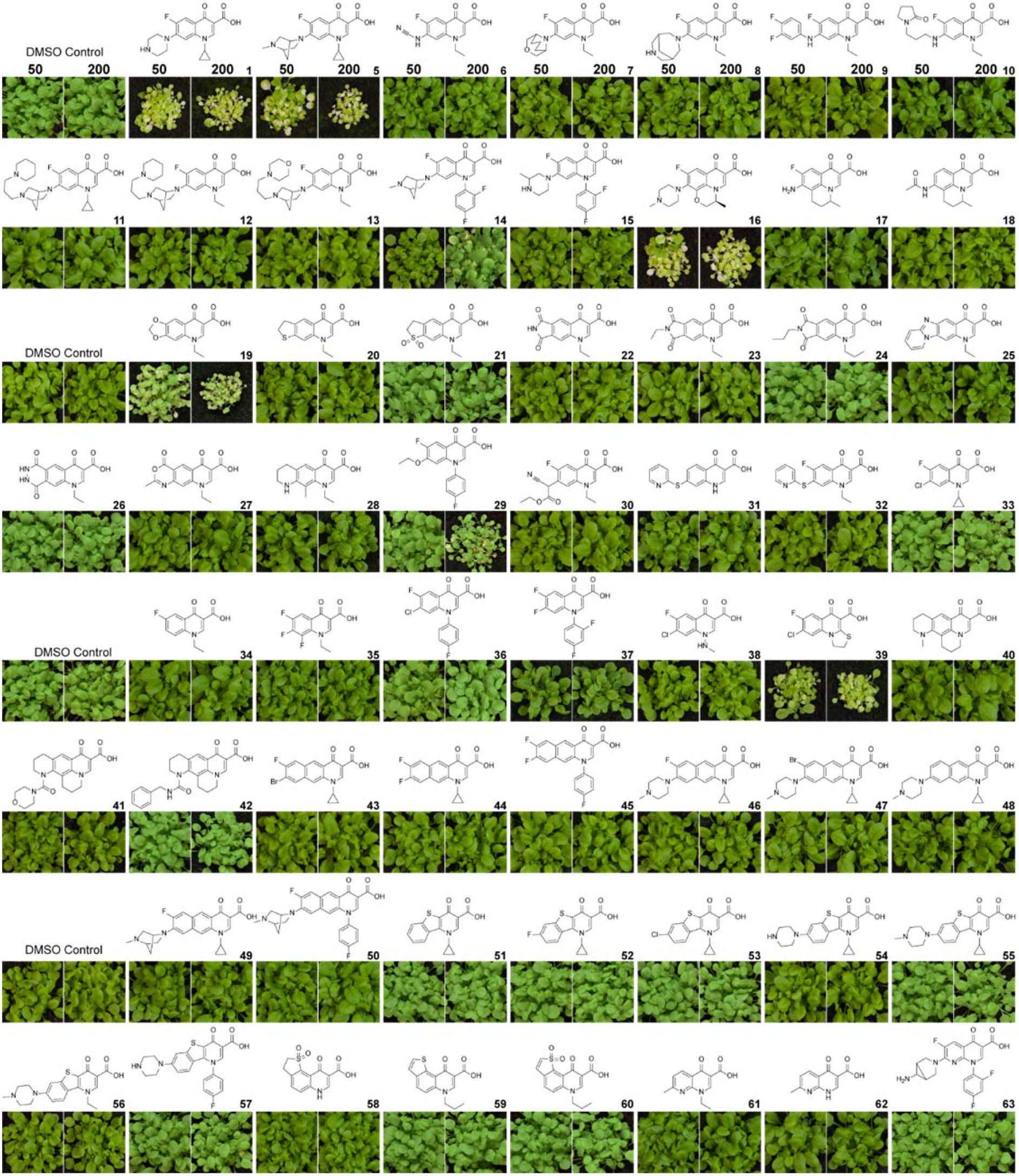
Herbicidal activity of ciprofloxacin analogue library. Wild type *A. thaliana* was grown on soil and treated with the relevant compound three- and six-days after germination. Each application was 0.5 mL of compound at 50 or 200 mg L^-1^, containing 2% DMSO and 0.02% Brushwet. Photos taken after 18 days of growth.

### Herbicidal potency of active compounds and their antibacterial activity

Compounds that possessed herbicidal activity were tested at a range of concentrations to further evaluate their potency (IC_50_). Levofloxacin **16** and the thiazolidine **39** had equivalent activity to **1** (Table 1). A slight reduction in activity was seen for danofloxacin **5** and oxolinic acid **19**. The 7-ethoxy derivative **29** had a significant reduction with only minor herbicidal activity.

**Table 1.** Herbicidal and antibacterial activity of herbicidal compounds. Herbicidal potency (IC_50_, mg L^-1^) of active compounds against *A. thaliana*, errors are expressed as standard error. Minimum inhibitory concentrations (MIC, mg L^-1^) against *E. coli* (K12), *P. aeruginosa* (ATCC 19429) and *S. aureus* (ATCC 25923).

| Compound | $IC_{50}$ ($mg\ L^{-1}$) | | MIC ($mg\ L^{-1}$) | |
| --- | --- | --- | --- | --- |
|  | *A. thaliana* | *E. coli* | *P. aeruginosa* | *S. aureus* |
| **1** | $11.1 \pm 1.8$ | 0.03125 | 0.125 | 1 |
| **5** | $23.4 \pm 3.4$ | 0.625 | 0.5 | 0.25 |
| **16** | $12.3 \pm 1.9$ | 0.625 | 0.5 | 0.25 |
| **19** | $22.9 \pm 3.8$ | 0.125 | 0.5 | 0.5 |
| **29** | $110 \pm 25$ | 32 | >128 | 32 |
| **39** | $13.6 \pm 1.3$ | 1 | 128 | 1 |

With these data in hand, the active analogues were tested for antibacterial activity against the model Gram-negative bacteria *Escherichia coli* and *Pseudomonas aeruginosa* and the Gram-positive bacterium *Staphylococcus aureus* (Table 1). All compounds tested had a lower activity against the Gram-negative bacteria than **1**, with clinical antibiotics **5, 16**, and **19** having only a slight decrease in MIC. The modestly herbicidal compound **29** had weak antibacterial activity. Interestingly, **39** which had the same herbicidal activity as **1** also had the same, albiet weak, antibacterial activity against *S. aureus* but a 32- and 1000-fold reduction in activity against the Gram-negative species *E. coli* and *P. aeruginosa*, respectively. Therefore, **39** demonstrates a scaffold with good herbicidal activity and weak antibacterial activity against both Gram-positive and Gram-negative bacteria, presenting a promising scaffold for further investigations.

In conclusion, from the results obtained here and seen previously, most structural changes to ciprofloxacin **1** severely diminish herbicidal activity. However, it has been demonstrated here that the fluoride at C-6 and a cyclic amino group at C-7 were not crucial for activity. Also, a phenyl group at N-1 was tolerated with respect to herbicidal activity, but there was only one example and generally a sterically small and mostly rigid motif is preferred in this region of the scaffold. Overall, further insight into scaffold design targeting this novel herbicidal mode of action would be greatly advanced with X-ray crystallographic studies of **1** with plant DNA gyrase.

## Acknowledgements

The authors thank Prof. Ulrich Jordis from Vienna University of Technology for the supply of analogues. MDW is supported by a Research Training Program Scholarship provided by the Australian Federal Government and The University of Western Australia. KAS and JSM also thank the Australian Research Council for funding (DP190101048).

## Author Contributions

KAS and JSM designed the research. MDW, AWD and KVS performed the research. MDW analysed the data. MDW, KAS and JSM wrote and edited the paper. All authors read and approved the final manuscript.

